# LPMX: A pure rootless composable container system

**DOI:** 10.1101/2021.06.04.445363

**Authors:** Xu Yang, Masahiro Kasahara

## Abstract

**Summary:** Delivering tools for genome analysis to users is often difficult given their complex dependencies and conflicts. Container virtualization systems such as Singularity isolate environments, helping developers avoid conflicts between tools. However, they lack *composability*, an easy way to integrate multiple tools in different containers or multiple tools both in a container and a host, which compromises the use of container systems in genome research. Another issue is that one may not be able to use a single container system of the same version at all sites they use, which discourages the use of container systems. To this end, we present a pure rootless composable container system, LPMX, that provides composability for letting developers easily integrate tools in different existing containers or on host, allowing researchers to compose existing containers. LPMX is pure rootless, so it does not require root privilege neither during installation nor at runtime, allowing researchers to use LPMX across sites without asking permissions from administrators. LPMX provides a pure userspace layered filesystem with at least an order of magnitude lower overhead for launching a new process than existing container systems. LPMX can import Docker and Singularity images.

**Availability and Implementation:** The source code of LPMX is available at https://github.com/jasonyangshadow/lpmx under Apache 2.0 License.

**Supplementary information:** Supplementary data are available at *Bioinformatics*online.

## 1 Introduction

Genome analysis usually involves shepherding data files through a bunch of tools and scripts, called pipeline or workflow (Leipzig, 2017; Koboldt *et al*., 2012; Lai *et al*., 2016; Kim *et al*., 2018). More complex genome analysis becomes, more third-party packages and tools are needed, and more likely there exist conflicting packages including different versions of the same package. This is known as the dependency hell. Tools used in genome analysis require varieties of environments such as dependent libraries or compiler versions. Setting up environments often costs a long time for researchers. This hampers researchers from using hard-to-install programs regardless of their scientific merits, making the progress of genome science significantly slower than it should be. The dependency hell is an urgent problem to address in genomics.

The community efforts, such as BioConda (Grüning *et al*., 2018), virtually eliminated the software installation problem for users, which is an enormous achievement. However, BioConda moved the burden of solving the dependency hell from users to developers, not having eliminated the problem itself. It is still developers’ responsibility to avoid conflicts between packages in BioConda and to create recipes for dependent libraries and programs. Besides, BioConda does not guarantee that any pairs of programs can be installed without conflicts within a single namespace; for example, Manta (Chen *et al*., 2016) requires Python 2, and installing any other tools requiring Python 3 in the same namespace will be conflicting. We can mitigate this issue by separating the conda namespaces. Suppose that tool X and Y conflict in a single namespace, and that you create a genome analysis pipeline Z that use tool X and Y. You wish that installing pipeline Z by conda would automatically install tool X and Y as well; however, this is impossible with the current conda system. We still need to eliminate the dependency hell for developers as well; container system isolates every single tool, making it easier to develop a pipeline software package that orchestrates many tools.

Container-based virtualization such as Singularity (Kurtzer *et al*., 2017), Docker (The Docker Community, 2020) or udocker (Gomes *et al*., 2018) enables developers to use their favorite Linux distributions and versions without struggling to make a program compatible with the BioConda build environment. Virtualization helps developers easily avoid the dependency hell issues during packaging. However, some types of pipelines are difficult to containerize often because containers are too isolated to integrate, which we call the lack of *composability*. For example, the Canu assembler (Koren *et al*., 2017) is difficult to package using current container systems because Canu inside container cannot submit a job to the batch job engine, which is outside container, in a straightforward manner. We need to *compose* the container and the host environment (the batch job engine). In contrast, BioConda does not suffer from such issues because tools and the host environments can interact directly with each other. Let us give another example. Suppose that a researcher develops a pipeline composed of several tools. They want to make these tools replaceable so users can dynamically plug these tools of different versions in for comparison. We want to make the pipeline container and the tool containers *composable* so we can test various combinations of the tools using the single pipeline. A new container system with composability is demanded.

Another issues with popular container systems in HPC such as Singularity are (1) users cannot create a new container image on HPC because they do not have a root privilege, (2) users hesitate to containerize small genomics tools, because a Singularity image must contain a full set of OS image even if the target tool is very small in size, and (3) when users have an account on multiple HPC sites, some sites completely lack container systems, or other sites may have Singularity but with different major versions, which makes them create many Singularity images even for a single tool. Pure rootless container engines such as udocker solves problem (1) and (3), while container engines that support layered file system (e.g., Docker) solves problem (2). However, none of the existing systems solve all of problems simultaneously.

Here, we propose a new rootless container engine, LPMX, that solves all these issues. LPMX provides composability for letting a container interface with the host or with another container. LPMX runs without a root privilege during runtime nor during installation, providing users with full features at any Linux clusters without administrators’ approval. LPMX has the first layered file system implemented purely in userspace, unlike an existing similar file system, FUSE-overlayfs(Open Repository for Container Tools, 2020); mounting a FUSE-overlayfs requires a root privilege under the default configuration of major distributions such RHEL 7/8, Ubuntu 19 or later.

## 2 Methods

The primary goals of LPMX are; a) To provide composability over containers and host; b) Being pure rootless; c) To support a layered file system.

To create a pure userspace container system, we developed a fake chroot environment based on fakechroot (dex4er, 2020), a project giving a chroot-like environment to end-users (non-root) by employing the LD_PRELOAD hack. The LD_PRELOAD hack enables us to inject arbitrary functions in dynamic libraries. By injecting special wrapper functions, tools see a fake virtual file system that does not really exist. For example, when a tool tries to open a file, an injected function replaces the path (in container) provided by the tool with a path to the real file on host, enabling processes in container open files in container without using root privilege. Using this technique, we developed a pure userspace layered file system, Userspace Union File System (UUFS), and integrated it into LPMX. The underlying data structure of UUFS is compatible with Docker, so we can easily import Docker images while retaining layers.

The composability is implemented in LPMX by wrapping exec^*^ functions in the GNU C library with the LD_PRELOAD hack. When a process inside container calls an exec^*^ function, LPMX traps it, and if the callee is one of the list of executables to compose, LPMX redirects the exec call to the target executable, which might be on host or in another container.

LPMX also has a specific General-Purpose Graphics Processing Unit (GPGPU) support as Docker, Singularity, and udocker do. A feature comparison Table 1 lists the key features of LPMX compared to existing popular implementations.

**Table 1.** Feature comparison between major container systems and virtual machines (VM). Y indicates ‘support’, N indicates ‘not support’.

| Features | LPMX | Docker | Singularity | VM | udocker |
| --- | --- | --- | --- | --- | --- |
| Pure rootless | Y | N | N | N | Y |
| Use Docker image | Y | Y | Y | N | Y |
| Use Singularity image | Y | N | Y | N | N |
| Support layered filesystem | Y | Y | N | N | N |
| Run statically linked programs | N | Y | Y | Y | Y |
| Composability | Y | N | N | N | N |
| GPGPU support | Y | Y | Y | Y | Y |

## 3 Results

To demonstrate the composability of LPMX, we performed two experiments that containerized the Canu assembler and composed different tools across containers (see Section 1 and 2 of *Supplementary Material*). Composing processes both on host and/or in containers with existing container engines is impossible without writing a significant amount of glue code.

UUFS is implemented in pure userspace. UUFS does not require an initial setup for a new process, so the time for launching a new process is reduced by up to 6 folds. To demonstrate the fast launching time of UUFS file system and a container in LPMX, we performed an experiment comparing container creation and destruction speed (see Section 3 of *Supplementary Material*).

One of the nice features with UUFS is that read()/write(), which are most frequently called functions in big data processing, do not have to be trapped; those two functions run at their native speed. To demonstrate that there is no performance overhead for calling untrapped functions such as read()/write() in LPMX, we performed an experiment that does structural variant calling analysis on human genome data at the SHIROKANE supercomputer. As expected, we observed no observable performance overhead in the total runtime (see Section 4 of *Supplementary Material*). On the other hand, calling trapped functions (such as open()) so many times would incur some penalty; we anticipate that a process in LPMX container will slow down to some extent when it calls trapped functions frequently. As an example of such worst-case for LPMX (with UUFS), we show an experiment in which we install and compile software packages. Compiling from source code and installing generated binaries involve very frequent metadata operations, which are all trapped by LPMX. LPMX with a few layers ran up to 1.5 times slower than the fastest compared system. However, we usually see this level of overhead only during creating a new container image, i.e., only once per developing a new container. To our knowledge, there is no practically used genomics pipeline with such a huge number of metadata operations, and therefore we believe that this level of overhead is acceptable. Note that the performance overhead depends on the number of layers (see Section 5 of *Supplementary Material*), as anticipated.

LPMX has a special support for GPGPU. We ran Guppy basecaller using the host GPGPU resources inside LPMX container to process Nanopore signals. As expected, we observed no observable performance overhead (see Section 6 of *Supplementary Material*).

To demonstrate how UUFS eases importing Docker images with multiple layers, we measured the required time to import images by Singularity and LPMX. LPMX was 2.5 times faster than Singularity (see Section 7 of *Supplementary Material*), presumably because Singularity has to merge all layers into one.

Last, the necessity of the rootless container system on High-Performance Computing (HPC) systems is discussed in Section 8 of *Supplementary Material*. The rootless nature of LPMX ensures that no additional security risk is ever introduced to any production environments (see Section 9 of *Supplementary Material*).

## 4 Conclusions

We developed LPMX, an open-source pure rootless composable container system that provides composability for allowing users to easily integrate tools from different containers or even from the host. LPMX accelerates the science by letting researchers compose existing containers and by letting researchers containerize tools/pipelines that are difficult to containerize using conda, saving the precious time of researchers. The technique used in LPMX allows it to run purely in userspace without root even during installation, ensuring that we can use LPMX at any Linux clusters with major distributions. The lowest overhead for launching containers with LPMX gives us courage to isolate tools as much as possible into small containers, minimizing the chance of conflicts. The support for the layered file system keeps the total size of container images for a single genomic pipeline modest, as opposed to Singularity that mostly uses a flat single-layer image. LPMX not only removes the burden from users but also from developers/researchers, saving the time of researchers, and thus accelerating the science.

## Acknowledgements

The super-computing resource was provided in part by Human Genome Center (the University of Tokyo), in part by NIG supercomputer at ROIS National Institute of Genetics.

