## Supplemental Material for "LPMX: A pure rootless composable container system"

### Supplementary Material

#### 1. CANU ASSEMBLER CONTAINERIZATION

Canu [1] assembler is a popular tool used in genome analysis research, which can assemble complete microbial genomes as well as eukaryotic chromosomes using either PacBio or Oxford Nanopore technologies. Canu is different from other analysis tools in a way that it can automatically detect and configure itself to use on most batch job systems, e.g. Univa Grid Engine (UGE), it can query the batch engine system and generate configuration scripts for job submission, and then submit these scripts to the batch engine system for distributed execution.

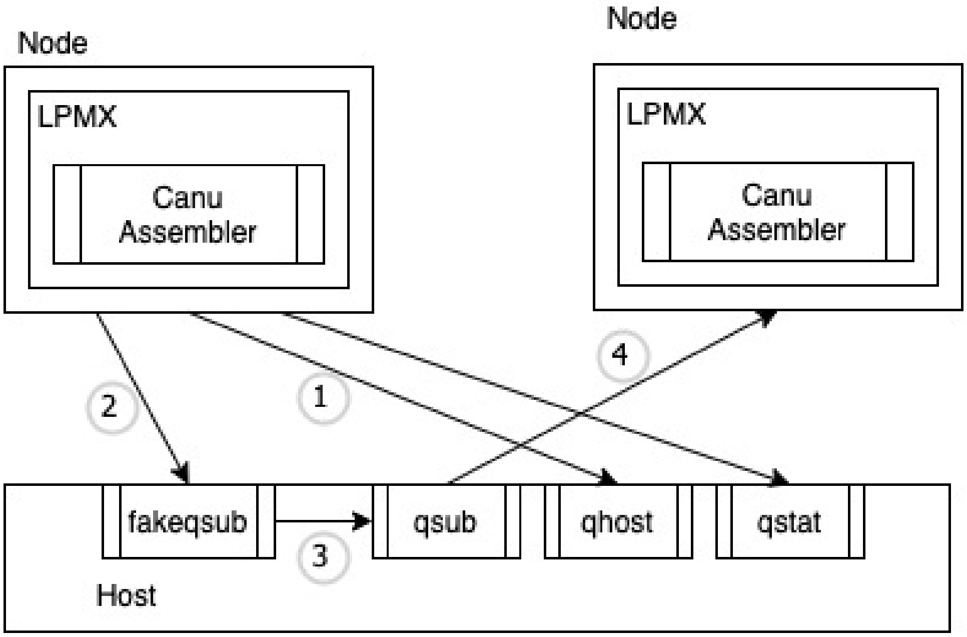


**Fig. S1.** An overview of the experiment, (1) By using LPMX, Canu assembler can easily call batch job commands (e.g. qhost) to query information from the job engine system. (2) When submitting a job (e.g. use qsub), the command is trapped and replaced by "fakeqsub", which is a delegate program that generates a shell script to restore containers on other compute nodes. (3) Then the "fakeqsub" calls the real qsub to submit the generated shell script. (4) When the generated shell script is executed on other compute nodes, the LPMX container is resumed and the containerized Canu assembler is launched for the analysis.

Traditional container systems cannot be used to containerize the Canu assembler without modifying files inside images, as these systems lack composability (i.e. do not allow the Canu assembler process inside containers to make a direct call to batch job engine commands on the host). LPMX readily solves this issue by featuring composability. LPMX provides executable the ability to call other executable on the host without modifying files inside containers so that batch job engine commands are visible to the Canu assembler inside containers. To handle the job submission for the Canu assembler, we developed a fake job wrapper running outside of containers to replace the default job submission command and delegate the action of resuming the container on the compute node. Fig. S1. shows that containerized Canu assembler can readily make a direct call to an executable on the host, and that the Canu remains fully workable inside LPMX thanks to the composability feature. We use *E. coli* K12 Oxford Nanopore dataset [1] with the Canu assembler version 2.1.1 on the supercomputer SHIROKANE [2] in the Human Genome

[1] http://nanopore.s3.climb.ac.uk/MAP006-PCR-1_2D_pass.fasta
[2] https://supcom.hgc.jp/shirokane.html

Center (HGC) equipped with UGE, to demonstrate the composability feature of LPMX.

The Canu assembler is containerized and works without modifying files; end-users can compose existing containers without modifying them.

#### 2. MULTIPLE APPLICATION VERSIONS TEST

In genome analysis exploratory experiments, combinations of different tools, as well as different versions of the same tools, are tested to get the best accuracy. Traditionally, researchers have to spend a large amount of time and effort containerizing all combinations manually. LPMX allows different tools and different versions of the same tools to be composed and integrated into the current container without launching new ones.

To demonstrate the function, a structural variation calling experiment on dataset *E. coli* K12 MG1655 [3] consisted of using different tools was performed inside a virtual machine created by VirtualBox (employing Vagrant). The virtual machine file and experiment scripts are available on Vagrant Cloud [2]. In this experiment, we injected different containerized tools, including bwa [3], minimap2 [4], samtools [5], and different versions of Genome Analysis Toolkit (GATK) [6], into a running container, which then dynamically composed these tools into different experimental trials. The experiment showed that LPMX allowed end-users to decouple and containerize tools into different containers while still made these containers cooperate to perform the exact same and complete analysis. Figure S2 shows an overview of the structural variation calling experiment in which different tools and different versions of the same tool were employed.

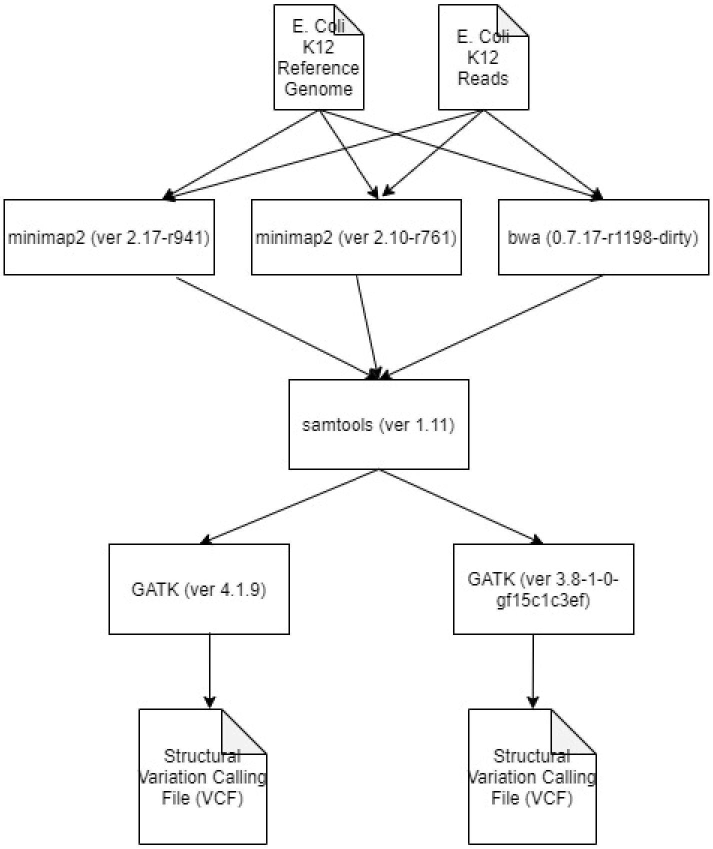


**Fig. S2.** An overview of the structural variation calling experiment. The input files are *E. Coli* K12 reference genome and reads, tools such as minimap2, bwa, samtools as well as GATK are employed to do the structural variation calling analysis.

[3] https://www.ncbi.nlm.nih.gov/genome/167?genome_assembly_id=161521, ftp://webdata:webdata@ussd-ftp.illumina.com/Data/SequencingRuns/MG1655/MiSeq_Ecoli_MG1655_110721_PF_R1.fastq.gz, ftp://webdata:webdata@ussd-ftp.illumina.com/Data/SequencingRuns/MG1655/MiSeq_Ecoli_MG1655_110721_PF_R2.fastq.gz

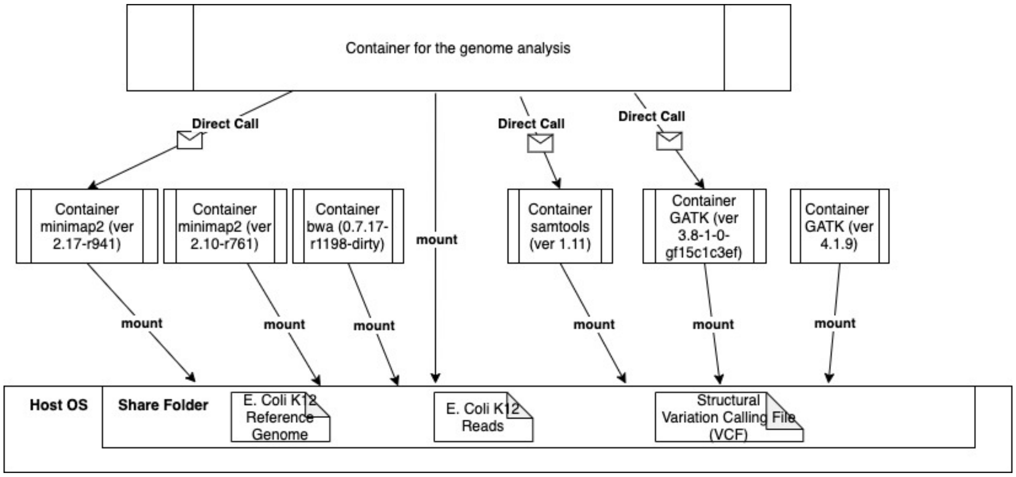


**Fig. S3.** An overview of the experiment. Various tools and different versions of the same tools were containerized into different containers that mount and share the same folder of the host. By exploiting the composability feature of LPMX, a container running genome analysis can make a direct call to exposed tools in other containers.

Figure S3 shows the overview of this experiment. We first separated and containerized different tools into different containers, but all containers shared the same directory in the host to allow data access and exchange. We pre-downloaded *E. coli* K12 dataset into the shared folder so that all containers could access it. Then we created a clean container (without any analysis tools installed), by using the composability feature, the newly created container can directly call existing tools as if they were installed locally and could complete the analysis using different combinations of these tools to achieve better experiment results without installing these combinations of tools into different containers, which saves a large amount of time and effort.

Thus, users can readily compose existing containers when building a large pipeline container without modifying files inside these containers.

#### 3. CONTAINER CREATION AND DESTRUCTION SPEED BENCHMARK

The Userspace Union File System (UUFS) inside LPMX supports a user-land layered filesystem so that containers sharing the same base image do not need to be re-created from scratch, greatly improving the container creation speed and saving local storage. Also, all binaries and libraries inside base images are no longer needed to be patched for preventing escape, which will take additional time for the first run and have to be restored when creating portable images. To compare the container creation and destruction speed of LPMX to those of Docker, Singularity, udocker [7], and Podman [8], we created a virtual machine using VirtualBox (employing Vagrant) and experimental scripts, which are available on Vagrant Cloud [9] and GitHub [10]. In this experiment, we created and destroyed the container ten times and measured the associated time cost using the shell-builtin (time) command. The (time) command measures the resource usage of a given running command and outputs the elapsed real time between invocation and termination, the user CPU time, and the system CPU time. We summed the user and the system CPU times to obtain the total time required to run the userspace logic code and execute the kernel space system calls during the run. The experiment was repeated five times and the data was averaged to represent a more accurate result.

##### A. Configuration

A created virtual machine served as the experimental environment. All container systems were installed and configured as follows:

- Docker version 19.03.6
- Podman version 1.6.2
- Singularity version 3.7.0

- udocker version 1.1.4
- LPMX version alpha-1.6.3

The host was equipped with an Intel i7-8750H processors @2.2GHz (6 cores and 12 threads) and 32 GB RAM memory. The virtual machine was version 6.1. The virtualized host was Ubuntu 18.04.5 LTS (Bionic Beaver). The memory size was 1024 MB. The containerized Guest OS was also Ubuntu 18.04 LTS (Bionic Beaver) downloaded from DockerHub using the tag Ubuntu:18.04. A shell script was executed in the virtualized host OS to evaluate the performance. The script creates and then immediately destroys the container and measures time cost.

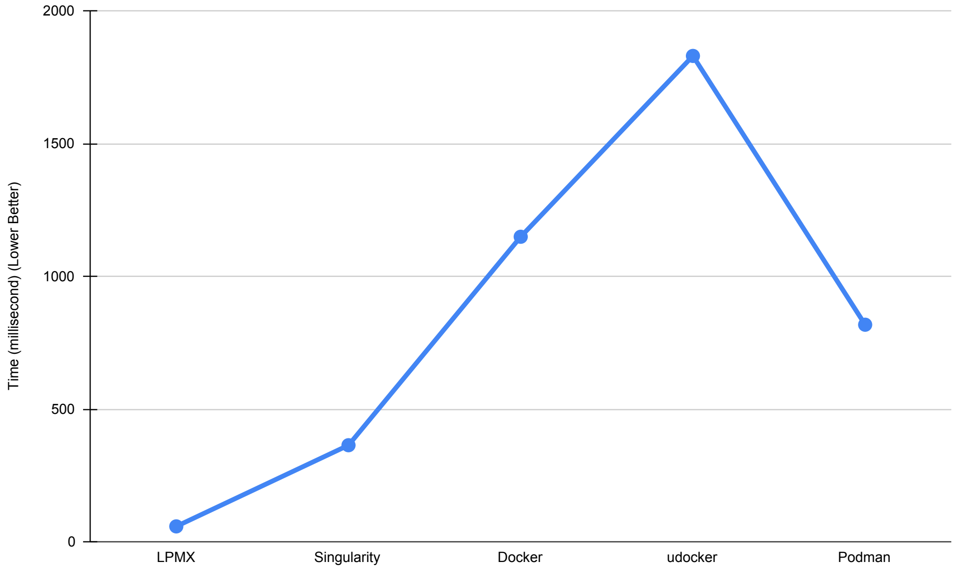


**Fig. S4.** The figure shows the average time cost (in ms; lower is better) required to create containers by different container systems, as shown in the figure, LPMX is the fastest, thanks to the Incorporated UUFS.

##### B. Result

As shown in Figure S4, LPMX is the fastest creator of container because the UUFS allows LPMX to create a read-write layer on top of existing read-only layers so that all base layers can be shared among containers running the same base image. As only a read-write folder and symbol links pointing to lower read-only layers are created, LPMX only requires little time to set up the container. udocker has to extract all layers and create a merged layer as the root file system each time when a container is launched, so more time is required.

Thus, by employing UUFS, LPMX can create and destroy containers much faster than other implementations, which makes LPMX suitable for running on HPC systems that often launch and destroy hundreds of containers simultaneously.

#### 4. STRUCTURAL VARIANTS CALLING EXPERIMENT ON SHIROKANE

To demonstrate that LPMX works on real scientific infrastructures and does not exhibit heavy performance overhead issues during genome analysis, we evaluated LPMX on the supercomputer SHIROKANE in the Human Genome Center located in the institute of medical science, the University of Tokyo. The LD_PRELOAD hack substantially wraps the original default functions and executes extra code in the userspace, which might trigger additional system calls. File system manipulation related operations will normally exhibit performance overhead. However, the read/write functions are kept untouched inside LPMX, which are common triggered in genome analysis, will not be slowed down. We performed a structural variant calling analysis on the GRCh38 human genome reads to show the performance overhead between execution on the bare host and in LPMX.We ran the experiments five times and calculated the averages.

##### A. Configuration

First, we installed the LPMX tool on the HGC supercomputer. Then we created different containers for tools used in this experiment. These tools are:

- samtools Version: 1.11 with htslib Version: 1.11
- minimap2 Version: 2.17-r941
- The Genome Analysis Toolkit (GATK) Version: 4.1.9.0-SNAPSHOT with HTSJDK Version: 2.23.0 Picard Version: 2.23.3

All data files were downloaded from the UCSC Genome Browser [4], and the reads files were downloaded from the international genome organization website [5]. We first used minimap2 to align the reads against the reference genome, then used samtools to process and polish the generated sam file, at last, used GATK toolkit to do structural variants calling analysis. We used the shell-builtin *time* command to evaluate the time cost for different steps. As the supercomputer SHIROKANE uses UGE as its batch job engine, for all experiments, we set the parallel environment option (pe) to def_slot with value 10, and set the highest virtual memory option (h_vmem) with the value 4 GB. The executions were identical using the bare host and LPMX.

##### B. Result

For minimap2 alignment step, both experiments mapped 10,640,477 sequences, the following Table S1 shows the performance returned by the minimap2 tool:

| Host | Real time (second) | CPU time (second) | Peek RSS (GB) |
|---|---|---|---|
| Bare Host | 406.343 | 1131.593 | 13.318 |
| LPMX | 391.447 | 1091.747 | 13.357 |

**Table S1.** Running the same experiment using Minimap2 with the same dataset on the bare host and LPMX. The real-time, CPU time and peek Resident Set Size (RSS) were recorded at the end of the experiment.

The samtools step involves many small processing steps, for example, creation a dict file for genome data and conversion of the data file from one format to another format, which lacks data statistic stability. Therefore, we skip the comparison of samtools. For the GATK, we used the output of samtools as the input of GATK and then evaluated the time cost in HaplotypeCaller, which is the core step of the structural variation calling and involves frequent read and write operations. The following Table S2 shows the results:

| Host | Processed Region | Time (minutes) |
|---|---|---|
| Bare Host | 11,717,231 | 40.99 |
| LPMX | 11,717,231 | 39.65 |

**Table S2.** GATK HaplotypeCaller time costs. "Processed" region indicates the number of processed regions in the genome. "Time" indicates the time cost during processing.

As the table shows, there is no obvious performance overhead in the real experiment. Read/write functions frequently called in genome analysis pipelines remain untouched in LPMX and thus are not slowed.

#### 5. PACKAGE INSTALLATION AND COMPILING OVERHEAD BENCHMARK

Here, we designed a performance benchmark test to reveal the worst performance overhead case of LPMX to other implementations, such as Docker, Singularity, Podman, and udocker. The

[4]https://hgdownload.soe.ucsc.edu/goldenPath/hg38/bigZips/
[5]https://www.internationalgenome.org/data-portal/sample/HG00171

experiment was designed and executed inside a virtual machine created by VirtualBox (employing Vagrant). The virtual machine file and experimental scripts are available on Vagrant Cloud [9] and GitHub [10]. We focused on file input/output (I/O) because pure calculation without any file I/O remains at the same speed by design. As the LD_PRELOAD hack substantially wraps original default functions and executes extra customized userspace code instead, which might trigger additional system calls compared to original direct calls. As software installation imposes a heavy I/O in genome analysis, we measured the overhead thereof. The root reason of performance overhead is caused by the execution of userspace logic code and extra system calls in kernel space. To evaluate the performance overhead of LPMX, common software and packages such as GCC, Python3, Ruby, Java Runtime Environment (JRE), and etc are installed and the most popular genome analysis tools as well as its dependencies, i.e. samtools and htslib, are compiled. The experiments were repeated five times and the average time costs measured by built-in *time* command were calculated. The result showed that compared to other container system implementations, performance overhead was observed up to 1.5 times in installing and compiling packages. However, users usually only need to install software once inside containers, meaning that the performance overhead is only suffered once. But for actual analysis repeatedly executed inside containers, only minimal performance overhead was observed as shown in Table S2. The reason is that functions, e.g. read/write, frequently called in the analysis are unwrapped in LPMX so that the raw performances are preserved.

We used the same environment used in section 3, and hence they shared the same machine specification. The experiment shell script was executed in the virtualized host OS to evaluate the performance. The two different tests in the script were:

1. Installing common packages via the system package manager, i.e apt.
2. Compiling popular software in genomics field, i.e samtools, and its dependencies including htslib.

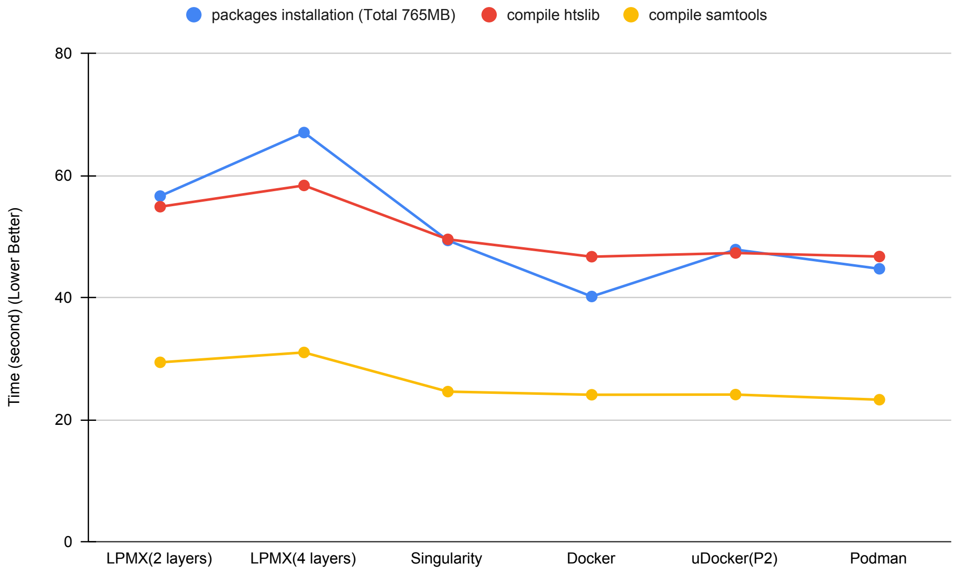


**Fig. S5.** Average time when installing and compiling the same packages in different container systems. Typically, LPMX exhibits a greater performance overhead compared with other implementations. Reduction of the search layer depth would reduce the overhead.

#### A. Result

As shown in Figure S5, compared with other container system implementations, the LPMX time cost is greater due to additional system calls in LPMX. LPMX.4.Layer is an experiment that launches the container on top of four layers, while LPMX.2.Layer is an experiment with only two layers. The overall performance overhead of LPMX is affected by the number of file operations and the layer depth.

In daily researches, researches normally install packages only once, while for any following calls to installed tools, there will be little performance overhead. Thus, the performance overhead during package installation and compiling would be acceptable.

#### 6. GUPPY BASECALLER GPU EXPERIMENT

GPGPU is becoming more popular as more genomics applications are optimized appropriately. Docker, Singularity, and udocker support GPGPU. We thus added GPGPU support to LPMX. As LPMX can expose arbitrary files and directories on the host to a container, we exposed GPGPU related files (especially special devices) to containers. To test if GPGPU can be used inside containers, we ran Guppy, a proprietary basecaller for Oxford Nanopore sequencers that uses GPGPU, inside an LPMX container, and confirmed that the outputs and the processing speed were as expected.

##### A. Configuration

- Guppy Basecaller Version: 3.4.5+fb1fbfb
- Intel(R) Xeon(R) CPU X5650 @2.67 GHz with 24 cores
- GPU: GeForce GTX 1080
- RAM memory: 192 GB
- Driver info: NVIDIA-SMI: 440.64.00 / Driver Version: 440.64.00 / CUDA Version: 10.2

We used the Chip137 IVT NA12878 RNA dataset [6] that contains 313 fast5 split files with a total size of approximately 35 GB. The following table S3 shows the information returned by the Guppy basecaller. We executed the experiment for five times and calculated the average values.

| Host | Time (seconds) | # Samples Called | # Sample per Second |
|---|---|---|---|
| Bare Host | 1344.233 | 18,614,227,490 | 1.38475e+07 |
| LPMX | 1342.606 | 18,614,227,490 | 1.38643e+07 |

**Table S3.** Performance information returned by the Guppy basecaller at the end of the experiment (caller time, called samples and calling speed).

##### B. Result

As shown above, considering the reasonable experiment performance fluctuation, the processing speed was as expected. As system calls related to hardware devices access, e.g. ioctl, are not intercepted in LPMX, raw performance is preserved.

Thus, LPMX supports accessing host GPGPU resources inside containers readily, which will benefit researchers in their daily researches.

#### 7. SINGULARITY AND LPMX IMPORTING DOCKER SAVED TARBALLS BENCHMARK

Docker can produce a tarred file containing all layers and metadata, which can be imported by other container systems to reproduce environments easily without recreating everything from scratch. Singularity and LPMX can import Docker saved tarballs to recreate the containers by reading the metadata, importing layers, and establishing correct runtimes. Singularity requires additional steps to squash extracted layers to create a read-only Singularity image, while LPMX can utilize incorporated UUFS mentioned in section 3 to directly mount extracted layers from saved tarballs and is thus more efficient than Singularity especially when importing a large amount of Docker saved tarballs. We measured the import performances of Singularity and LPMX using the same environments mentioned in section 3. We randomly selected five Docker images from the Biocontainers repository on the Docker Hub, imported these images using

[6] https://s3.amazonaws.com/nanopore-human-wgs/rna/links/NA12878-IVT-RNA_All.files.txt

Singularity and LPMX, and measured the time costs using Linux built-in *time* command. We ran the experiment five times and calculated the average times. Compared with LPMX, Singularity was 2.5 times slower. By benefiting the UUFS system, LPMX can directly mount layers extracted from tarballs and launch containers on these layers without additional works.

#### 8. DISCUSSION ON THE NECESSITY OF PURE ROOTLESS CONTAINER SYSTEM ON HPC SYSTEMS

Rootless in container technology refers to the ability that unprivileged users can create and manage containers. Pure rootless defined in the manuscript refers to the ability that privileges are not required even when setting up the container system runtime, meaning that privileges are not asked in any stages of using the container system. This feature is desired because users can use the same container system at multiple sites regardless of Linux kernels and distros. Also, the attack surface of rapidly evolving software on the host is minimized and no additional potential risks will be introduced, which is suitable for multiple-tenant systems.

##### A. Security Concerns

When a researcher asks administrators to install Singularity [11] or Docker, which are the two most popular container-based virtualization systems, the administrators are concerned about the security. Docker, which uses a daemon with a root privilege, had several critical security bugs [12] in the past that might have led to a serious privacy breach in human genome analysis, which is one of the biggest applications in genome analysis. The recent Docker version implements a non-root mode [13] to reduce such concerns, but it still suffers security issues from using the unprivileged user namespace, which might let the container system be compromised by malicious code running inside unprivileged processes. The reason behind this security issue is that capabilities acquired by unprivileged processes in the new namespace might help these processes gain root privileges by using zero-day bugs in the Linux kernel. The example [7] belongs to such cases. Besides, to enable the non-root mode, necessary packages, e.g. newuidmap, fuse-overlayfs are needed to be installed in advance, which also requires root privilege and introduces new security risks. Singularity is considered as much more secure than Docker and therefore is preferable in the context of genome analysis, though users have to ask administrators to install Singularity if it is not available on the system. Singularity requires a root privilege to install, which puts a huge burden on users to persuade the administrators to actually install Singularity.

Our system, LPMX, does not require root in both during installation and runtime of containers. The pure rootless design also eliminates security concerns as it does not enhance any privileges in userspace.

##### B. Kernel Support

To support rootless container systems on HPC systems, a modern Linux kernel is usually needed. For example, to support unprivileged user namespace, the kernel version should be higher than version 3.18 with security patches, and additional modifications to the Linux kernel have to be made, e.g. manually enable the kernel option kernel.unprivileged_userns_clone. Except that there exist security concern of unprivileged user namespace in the previous section, to make rootless container system work properly, e.g. Docker non-root mode, it often requires the server administrators to upgrade the current stable Linux system to a more recent one, e.g. CentOS 7.7, so that other necessary packages, e.g. fuse-overlayfs, can be successfully installed. Upgrading the current system to a recent one is always good unless we afford to do so, but it usually means additional work has to be done before and after the system upgrade.

Our system, LPMX, is pure rootless in userspace and is robust to Linux kernel, and hence there is no need to modify the current Linux kernel of the host.

##### C. Vendor Support

Administrators of HPC systems sometimes refuse to install container systems due to contract issues with vendors. As discussed in the previous section, supporting non-root mode container systems usually involves upgrading Linux kennels for new features, this also means administrators have to get more support from contract vendors, which might cost a long time in public institutions. This becomes prominent especially when special hardware and drivers are used. For example, the Infiniband network driver is used on the supercomputer in the National Institute of

[7] https://www.openwall.com/lists/oss-security/2020/09/03/3

Genetics (NIG) in Japan, to install and support Singularity, the driver has to be recompiled every time the kernel is updated, which requires extra work and effort.

Our system, LPMX, does not require a root privilege, and hence it does not require administrators' approval nor additional financial costs associated with support and maintenance.

#### 9. SECURITY AND LIMITATIONS

##### A. Security

In contrast with a Docker daemon that have potential security risks, LPMX is a pure rootless, non-setuid program. Therefore the use of LPMX will not increase any security risks by design. Any operations requiring a root privilege is not permitted. For example:

1. Access host protected devices and files
2. Listen on privileged ports (below 1024)
3. Mount file systems (on host)
4. Change system settings including userid, groupid, system time, network-related things such as firewall rules and interfaces.

##### B. Limitations

Some limitations are listed as followings, but not limited to:

1. Executables statically linked do not work properly inside containers. Recompiling them with shared libraries is a recommended workaround. Alternatively, users can install such statically linked executables on host and call it from inside container by exposing them by LPMX, if needed.
2. The more layers an image has, the heavier performance overhead end-users will see; merging many layers into one layer during the creation of containers can be a workaround.
3. Some commands, e.g ps command, will not work as expected inside containers due to the lack of inter-process communication namespace isolation; a customized ps command wrapper can do the trick.
4. LPMX does not work with a root account; end-users should use non-privileged accounts.
5. Setuid/setgid executables do not work inside LPMX containers because LD_PRELOAD is disabled by Linux for such executables.
6. When executables uses a system call that does not exist in the host kernel, LPMX cannot execute them. This is the common limitation of container systems.
7. As LPMX only wraps functions in the standard C library, i.e. glibc, LPMX will not run on other glibc independent Linux distros, e.g. Alpine Linux.

These limitations usually would not be a problem when users execute a genome analysis pipeline with only open-sourced tools that inputs files and outputs files without interacting with network sockets, which is the common case for genome analysis pipelines.
